# Heavy metal pollution characteristics of soil around a copper-nickel mine tailings pond in the northwest arid area of China and evaluation of desert phytoremediation potential

**DOI:** 10.1101/2022.10.24.513495

**Authors:** Jianfei Shi, Zhengzhong Jin, Zhibin Zhou, Xin Wang, Xiaoliang Yang, Wenting Qian

**Author notes:** Corresponding author: Wenting Qian.

## Abstract

Phytoextraction and phytostabilization are two mechanisms used by plants to remove and stabilize heavy metals in soil. However, there has been little research on the extraction/stabilization of heavy metals by desert plants in arid areas. This study analyzed eight heavy metals (Cr, Ni, Cu, Zn, Cd, Pb, Mn, and As) in 10 desert plants and their growth matrices. In addition, the single factor pollution index and Nemerow comprehensive pollution index were used to evaluate soil pollution. Based on the biological accumulation factor and the biological concentration factor, the fuzzy evaluation method was used to evaluate plant extraction/stability potentials for heavy metals. The results showed that Cd, Cu, Ni, and Cr in the soil around the tailings pond were at the heavy pollution level, Mn and Pb pollution was moderate, and Zn and As pollution was light. The matrix factorization model showed that Cu and Ni came from industrial pollution; Cd and Cr came from atmospheric deposition and agricultural pollution; Pb came from traffic pollution; and Mn, Zn, and As came from natural sources. The metal contents of the desert plants exceeded the standard for normal plants, of which Cr contents in the *Atriplex patens* and *Ammophila breviligulata* Fernald aboveground parts were 35.63 and 53.77 mg/kg respectively, and the Ni contents in the *Klasea centauroides* subsp. *polycephala* (Iljin) L. Martins underground parts and the *A. breviligulata* Fernald aboveground parts were 102.67 and 101.67 mg/kg, respectively, exceeding the maximum toxicity threshold for Cr and Ni. *Ammophila breviligulata* Fernald had the highest plant comprehensive extraction coefficient (CEI) and plant comprehensive stability coefficient (CSI) at 0.81 and 0.83 respectively, indicating that it has strong heavy metal extraction and stabilization abilities. Therefore, *A. breviligulata* Fernald can be selected as a remediation species for heavy metal pollution in the soil around tailings reservoir areas.

## 1. Introduction

Soil heavy metal pollution has gradually become a global problem due to increased industrialization and urbanization (Hanfi et al., 2020; Jeong et al., 2020; Khodaverdiloo et al., 2020; Yu et al., 2020). In particular, mining causes damage to regional vegetation (Li et al., 2020; Mwitwa et al., 2012) and its tailings from ore refining are an important source of environmental pollution (Consuelo et al., 2021; Zhang et al., 2021). Tailings are residual wastes from the processing and production of ores and industrial minerals. In particular, they contain unstable primary and secondary minerals (Huang et al., 2012). It is estimated that more than 10 billion tons of tailings are produced in the world every year and a large amount of tailings accumulate leading to a large environmental footprint in time and space (Adiansyah et al., 2015). Generally, tailings are acidic (Mendez et al., 2008), mainly composed of silt or sand sized particles, lack nutrient elements (N, P, K) to support plant growth, and contain almost no organic matter, but their heavy metal contents are high (Ginocchio, 2000; Mendez and Maier, 2008). In arid climates, the fine particles on the tailings surface are rich in pollutants and are vulnerable to the influence of wind-induced dispersion and hydraulic action. This means that they continue to diffuse to the surrounding environment, eventually leading to expansion of the polluted area (G et al., 2015; Mendez et al., 2008). In addition, the large evaporation rate in arid areas makes the water rapidly migrate upward through capillary action, providing supersaturation conditions for the formation of the secondary evaporation solid phase by weathering (Khorasanipour, 2015). Therefore, soil heavy metal pollution in arid areas may pose a greater threat to the ecological environment than in other areas, such as humid or semi-humid areas (Punia, 2021).

Heavy metal elements are not affected by biodegradation, but are converted between various species, which means that it is very difficult to remediate heavy metals in soil (Sarwar et al., 2017). In recent decades, researchers have tried to develop effective remediation technologies for contaminated soils (Khodaverdiloo et al., 2020). Currently, three general methods are used to remediate heavy metal pollution in soil. These are physical remediation, chemical remediation, and biological remediation (Roychowdhury et al., 2018). Physical remediation methods include soil replacement, soil washing, and vitrification, while soil chemical methods include stabilization and extraction (Ashraf et al., 2019; Roychowdhury et al., 2018). However, physical and chemical remediation methods are not suitable for large-scale application due to high cost, low efficiency, or possible damage to the soil structure (Khodaverdiloo et al., 2020; Zhao et al., 2014). In contrast, phytoremediation methods (bioremediation methods) have attracted extensive attention due to their advantages, such as being solar energy driven, their cost-effectiveness, and they are environmentally friendly (Sarwar et al., 2017). Therefore, evaluating the phytoremediation potential for heavy metals in heavy metal contaminated sites in arid areas is of great significance when attempting to enrich the database of heavy metal enriched plant resources and promote the wide applicability of heavy metal phytoremediation technology.

Phytoremediation technology refers to the various technologies that use natural plants or transgenic plants to repair polluted environments (Flathman et al., 1998). Phytoremediation techniques include plant extraction, plant stabilization, rhizosphere filtration, plant degradation, and plant volatilization (Karaca et al., 2018), while phytoextraction and plant stabilization techniques are usually used to remediate heavy metal pollution in soil (Mendez et al., 2008; Wong, 2003). Plant extraction absorbs and transfers heavy metal pollutants to the aboveground part through plant roots and finally removes the heavy metals in soil through artificial harvesting and treatment (Lam et al., 2018; Padmavathiamma et al., 2007). Plants also stably absorb and accumulate heavy metals in soil through their roots, which reduces heavy metal mobility and effectiveness and stabilizes (harmlessness) heavy metals in soil (Lam et al., 2018; Padmavathiamma et al., 2007). Based on these effects, this study evaluated the main role of phytoremediation technology in soil heavy metal remediation by using fuzzy theory and the plant comprehensive extraction coefficient (CEI) and plant comprehensive stability coefficient (CSI). The CEI and CSI for plants can reflect the removal and stabilization effect of plants on different heavy metals in soil and provide a more reasonable basis that can be used to select plants for the remediation of complex heavy metal contaminated sites.

In summary, the aims of this study were (1) to explore the impact of tailings accumulation on the enrichment of heavy metals in the surrounding soil, (2) clarify the sources of heavy metals in soil and provide reasonable suggestions for the effective prevention and control of heavy metal pollution in soil, and (3) evaluate the phytoremediation potential of native plants to provide a reference basis for the phytoremediation of heavy metal contaminated soil.

## 2. Materials and methods

### 2.1. Study area description and sample collection

The copper-nickel mine tailings pond was built in 1999 and is located in northern Xinjiang Province and in the eastern part of Fuyun County (Fig. 1). The design level of the pond capacity is grade Ⅴ and the floor area is about 1.5 × 10^5^ m^2^. The slag waste is disposed of by the traditional wet discharge method and the discharge volume is about 1,000 t/d. The altitude of the area ranges from 500–900 m and the soil types are mainly light brown calcium soil and typical brown calcium soil. The area is subjected to a continental cold temperate climate. It is dry and windy in spring, but cold in winter. There are also large temperature differences between day and night. The annual average temperature is 3.0°C, the annual precipitation is 186.4 mm, the annual evaporation is 1,829.7 mm, the annual extreme maximum temperature is 42.2°C, and the extreme minimum temperature is −51.5°C. It is one of the high, cold regions in China. In September, 2021, the natural plants around the tailings pond were investigated and sampled. During this survey, 10 native plants belonging to eight families were collected and recorded (Table 1). The plant samples were then divided into underground and aboveground parts and each plant was replicated three times. Soil samples from around the plant roots were collected at the same time as the plant samples. The soil sampling depth was 0–20 cm and a total of 60 plant samples and 30 soil samples were collected.

**Table 1.** Composition of native plant species around the tailing compounds.

| Species | Site | Family | Life form |
| --- | --- | --- | --- |
| *Polygonum aviculare* L. | 9 | Polygonaceae | Annual grass |
| *Salsola ruthenica* | 6 | Chenopodiaceae |  |
| *Halogeton arachnoideus* Moq. | 10 | Amaranthaceae |  |
| *Atriplex patens* | 8 | Chenopodiaceae |  |
| *Klasea centauroides* subsp. *polycephala* (Iljin) L. Martins | 4 | Compositae | Perennial grass |
| *Limonium sinense* (Girard) Kuntze | 1 | Plumbaginaceae |  |
| *Sphaerophysa salsula* (Pall.) DC. | 2 | Leguminosae |  |
| *Ammophila breviligulata* Fernald | 5 | Gramineae |  |
| *Phragmites australis* (Cav.) Trin. ex Steud | 3 | Gramineae |  |
| *Peganum harmala* L. | 7 | Zygophyllaceae |  |

**Figure 1.**
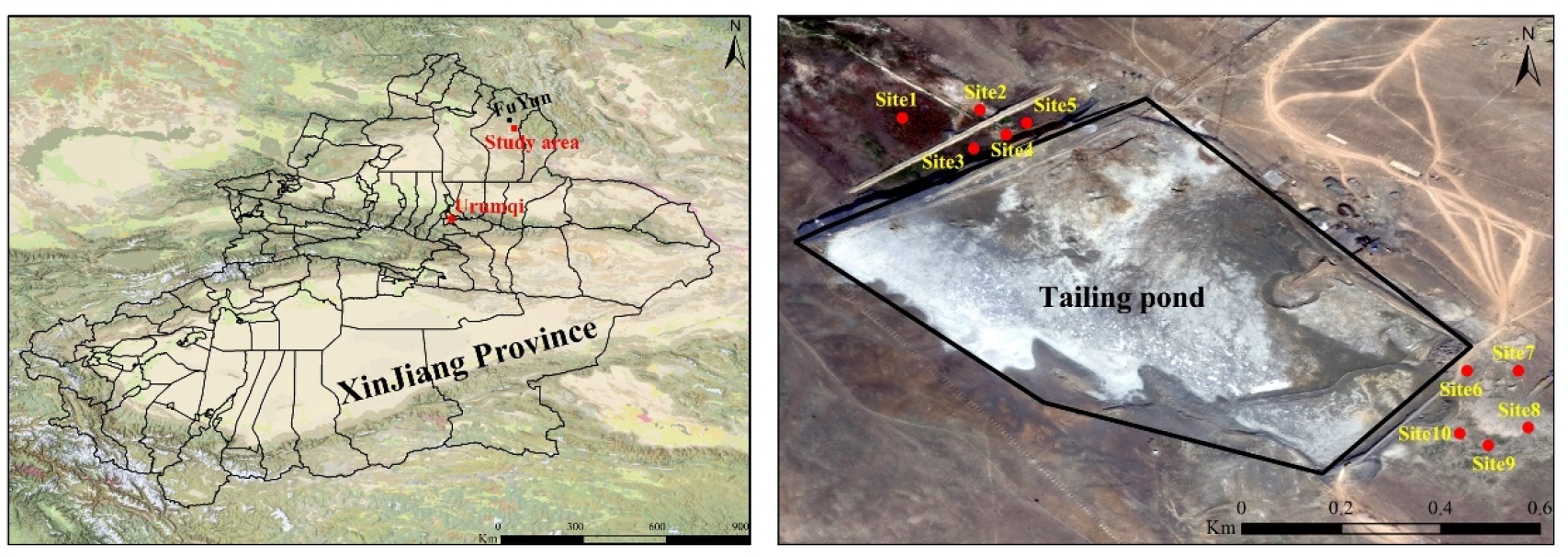
Location of the study area and distribution of the sample points.

### 2.2. Sample processing and determination

The plant samples were first washed with tap water and then three times with deionized water. They were placed in an oven at 105°C for 20 minutes, dried at 70°C to a constant weight, ground with a grinder to pass through a 100 mesh nylon sieve, and bagged. Then, 0.2 g subsamples of the plants were weighed out, placed in a microwave digester, and digested in an HNO_3_-H_2_O_2_ digestion system (HNO_3_:H_2_O_2_ = 5:1, volume ratio) until the liquid clarified. The soil samples were dried after removing impurities, such as stones and animal and plant residues. These samples were then ground with agate mortar through a 200 mesh nylon sieve, and bagged until needed. Then, 0.1 g subsamples of the soil were digested in a microwave digester under a HNO_3_-HF-HClO_4_ digestion system (HNO_3_:HF:HClO_4_ = 3:1:1, volume ratio) until the liquid clarified. The experiment was verified using the blank control method, the double parallel sample method, and the standard addition recovery method to ensure the accuracy of the experimental and determination processes. The supernatants from the subsamples were removed and the contents of eight heavy metals (Cr, Ni, Cu, Zn, Cd, Pb, Mn, As) were determined by inductively coupled plasma mass spectrometry (ICP-MS).

### 2.3. Evaluation of heavy metal pollution in soil

#### 2.3.1. Single factor pollution index method

The single factor pollution index (*P_i_*) is a common method used to evaluate the degree to which a soil has been polluted with heavy metals (Wang et al., 2011). The index is used to evaluate environmental quality by comparing the measured value with the standard value. The calculation formula is as follows (Wang et al., 2011; Young et al., 2021):

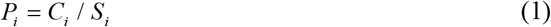

where *P_i_* is the single-component contamination index, *C_i_* is the measured concentration of examined metal *i* in the soil, and *S_i_* is the background concentration of metal *i*. This study took the background value for heavy metals in Xinjiang soil as the standard value. The evaluation results were divided into five grades: *P_i_* ≤ 0.7, safe; 0.7 < *P_i_* ≤ 1.0, warning; 1 < *P_i_* ≤ 2, slight pollution; 2 < *P_i_* ≤ 3, moderate pollution; and *P_i_* > 3, heavy pollution.

#### 2.3.2. Nemerow comprehensive pollution index method

The Nemerow comprehensive pollution index is used to calculate the comprehensive pollution effects of all assessed heavy metals. It can comprehensively reflect the effects of various heavy metals on soil and avoids the weakening of heavy metal weights caused by averaging. The calculation formula is as follows (Liang et al., 2011; Liu et al., 2020; Nemerow, 1991):

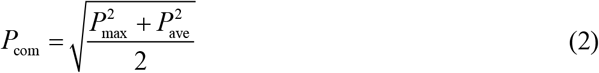

where *P*_com_ is the composite contamination index, *P*_ave_ is the average value of the single-factor index, and *P*_max_ is the maximum value of the single-factor index. The evaluation results were divided into five grades: *P*_com_ ≤ 0.7, safe; 0.7 < *P*_com_ ≤ 1.0, warning; 1 < *P*_com_ ≤ 2, light pollution; 2 < *P*_com_ ≤ 3, moderate pollution; and *P*_com_ > 3, heavy pollution.

### 2.4. Evaluation of phytoremediation potential

#### 2.4.1. Enrichment characteristics of heavy metals in plants

Heavy metals absorbed by plants are mainly characterized by the biological accumulation factor (BAF) and the biological concentration factor (BCF). Their calculation formulas are as follows (Elalaoui et al., 2021):

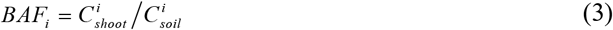

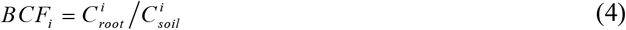

where *i* represents the ith heavy metal element; *BAF*_*i*_ is the enrichment coefficient of plants for the *ith* heavy metal, which is a parameter that is used to evaluate the ability of plant stems and leaves to extract heavy metals from soil; *BCF*_*i*_ is the stability coefficient of plants for the *ith* heavy metal and is a parameter used to evaluate the ability of plant roots to stabilize heavy metals; 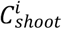 is the content of the *ith* heavy metal in the aboveground part of the plant; 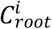 is the content of the *ith* heavy metal in the underground part of the plant; and 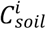 is the content of the *ith* heavy metal in the plant growth matrix.

#### 2.4.2. Comprehensive extraction index for plants

The comprehensive extraction index (CEI) for plants is based on fuzzy synthesis and can be used to evaluate the plant comprehensive extraction potential for various heavy metals under multiple heavy metal combined pollution conditions. The calculation formula is as follows (Hołtra et al., 2020; Zhao et al., 2014):

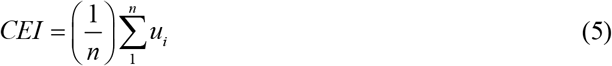

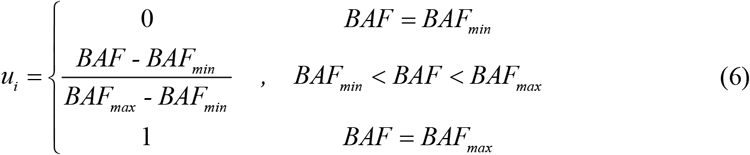

where *CEI* is the comprehensive extraction index for plants; *n* is the total number of heavy metals analyzed; *BAF* is the biological accumulation factor of plants for the *ith* heavy metal; *BAF*_*max*_ and *BAF*_*min*_ are the maximum and minimum values of the biological accumulation factor for *ith* heavy metals in the investigated plants, respectively; and *u*_*i*_ is the fuzzy membership value. The comprehensive extraction potential of plants is divided into three grades, poor (CEI ≤ 0.4), good (0.4 < CEI < 0.7), and excellent (CEI ≥ 0.7).

#### 2.4.3. Plant comprehensive stability index

The plant comprehensive stability index (CSI) is based on fuzzy synthesis and can be used to evaluate the plant root comprehensive stability potential for various heavy metals under multiple heavy metal combined pollution conditions. The calculation formula is as follows:

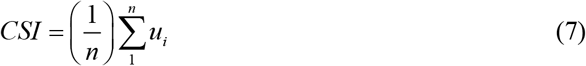

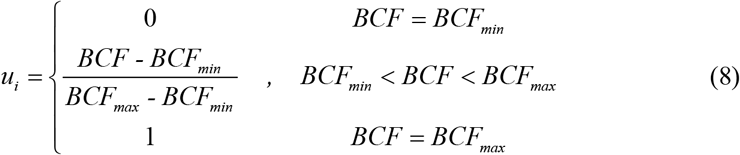

where *CSI* is the comprehensive stability index for a plant; *n* is the total number of heavy metals analyzed; *BCF* is the biological accumulation factor of plants for the *ith* heavy metal; *BCF*_*max*_ and *BCF*_*min*_ are the maximum and minimum values of the biological concentration factor for *ith* heavy metals in the investigated, plants, respectively; and *u*_*i*_ is the fuzzy membership value. The comprehensive stability potential of plants is divided into three grades, poor (CEI ≤ 0.4), good (0.4 < CEI < 0.7), and excellent (CEI ≥ 0.7).

### 2.5. Data processing and analysis

IBM SPSS statistics 22.0 (IBM Corp., Armonk, NY, USA) was used to statistically analyze the heavy metal contents in the soil, EPA PMF 5.0 was used for the source analysis of heavy metals in soil, and Origin 2020 (OriginLab Corporation, Northampton, MA, USA) and ArcGIS 10.6 (ESRI Inc., Redlands, CA, USA) were used for the graphics.

## 3. Results and analysis

### 3.1. Statistical characteristics of soil heavy metal content

According to the descriptive statistical analysis (Table 2), the Cr, Ni, Cu, Zn, Cd, Pb, Mn, and As contents in the soil samples were 255.2–911.3, 393.6–2861.1, 307.9–1654.3, 99.4–151.6, 2.7–12.7, 18.1–69.6, 1318.5–1810.8, and 5.4–4.9 mg/kg, with average contents of 529.0, 1168.0, 738.5, 118.2, 6.4, 40.1, 1600.3, and 13.9 mg/kg, respectively. The average Cr, Ni, Cu, Zn, Cd, Pb, Mn, and As contents in the soil samples were 10.7, 43.9, 27.7, 1.7, 53.3, 2.1, 2.3, and 1.2 times more than the background values for heavy metals in Xinjiang soils, respectively. Compared to the national soil environment class II standard, Cr, Ni, Cu, Cd, and As in the soil samples exceeded the standard by about 3.5, 29.2, 14.8, and 21.3 times, respectively. The variation indexes for Ni and Cu in the soil were 61.72% and 53.16% respectively, indicating that Ni and Cu are causing serious localized pollution. The variation indexes for Cr, Cd, Pb, and As were 47.19%, 37.94%, 43.25%, and 38.22%, respectively, which showed that the variation indexes were relatively weak. The variation indexes for Zn and Mn were 11.87% and 9.38%, respectively, which are less than 25% and shows that the variation was low and that they are less affected by external conditions.

**Table 2.** Statistics for heavy metal content in the soil around the tailing reservoir area (n = 30)

| Heavy metal | Cr | Ni | Cu | Zn | Cd | Pb | Mn | As |
| --- | --- | --- | --- | --- | --- | --- | --- | --- |
| Mean(mg/kg) | 529.0 | 1168.0 | 738.5 | 118.2 | 6.4 | 40.1 | 1600.3 | 13.9 |
| Minimum(mg/kg) | 255.2 | 393.6 | 307.9 | 99.4 | 2.7 | 18.1 | 1318.5 | 5.4 |
| Maximum(mg/kg) | 911.3 | 2861.1 | 1654.3 | 151.6 | 12.7 | 69.6 | 1810.8 | 24.9 |
| Coefficient of Variation (%) | 47.19 | 61.72 | 53.16 | 11.87 | 37.94 | 43.25 | 9.38 | 38.22 |
| BG<sub>1</sub><sup>a</sup> (mg/kg) | 49.3 | 26.6 | 26.7 | 68.8 | 0.12 | 19.4 | 688 | 11.2 |
| BG<sub>2</sub><sup>b</sup> (mg/kg) | 150 | 40 | 50 | 200 | 0.3 | 250 | - | 40 |

<sup>a</sup>BG<sub>1</sub>: background values for heavy metals in Xinjiang soil
<sup>b</sup>BG<sub>2</sub>: National Soil Environment class II standard

### 3.2. Evaluation of heavy metal pollution in soil

The pollution caused by different heavy metal elements in the soil was evaluated using Formula (1). The results (Fig. 2) show that the single factor pollution indexes for heavy metal elements Cd, Cu, Ni, Cr, Mn, Pb, Zn, and As were between 22.5–108.5, 11.5–62.0, 14.8–107.6, 1.7–6.1, 1.9–2.6, 0.9–3.6, 1.4–2.2, and 0.5–2.2, respectively. Among them, the average single factor pollution indexes for Cd, Cu, Ni, and Cr were 53.2, 27.7, 43.9, and 3.5, respectively, which were greater than 3 and meant that they reached the heavy pollution grade; the single factor pollution index mean values for Mn and Pb were 2.3 and 2.1, respectively, which were at the moderate pollution level; and the single factor pollution index mean values for Zn and As were 1.7 and 1.2, respectively, which were at the light pollution level. The Nemerow comprehensive pollution index results, calculated using Formula (2) (Fig. 2), showed that the Nemerow pollution indexes for all sample points ranged from 25.0 to 79.0, with an average of 43.9, which was at the heavy pollution level. In general, the soil was heavily polluted with metals and remediation measures were urgently needed.

**Figure 2.**
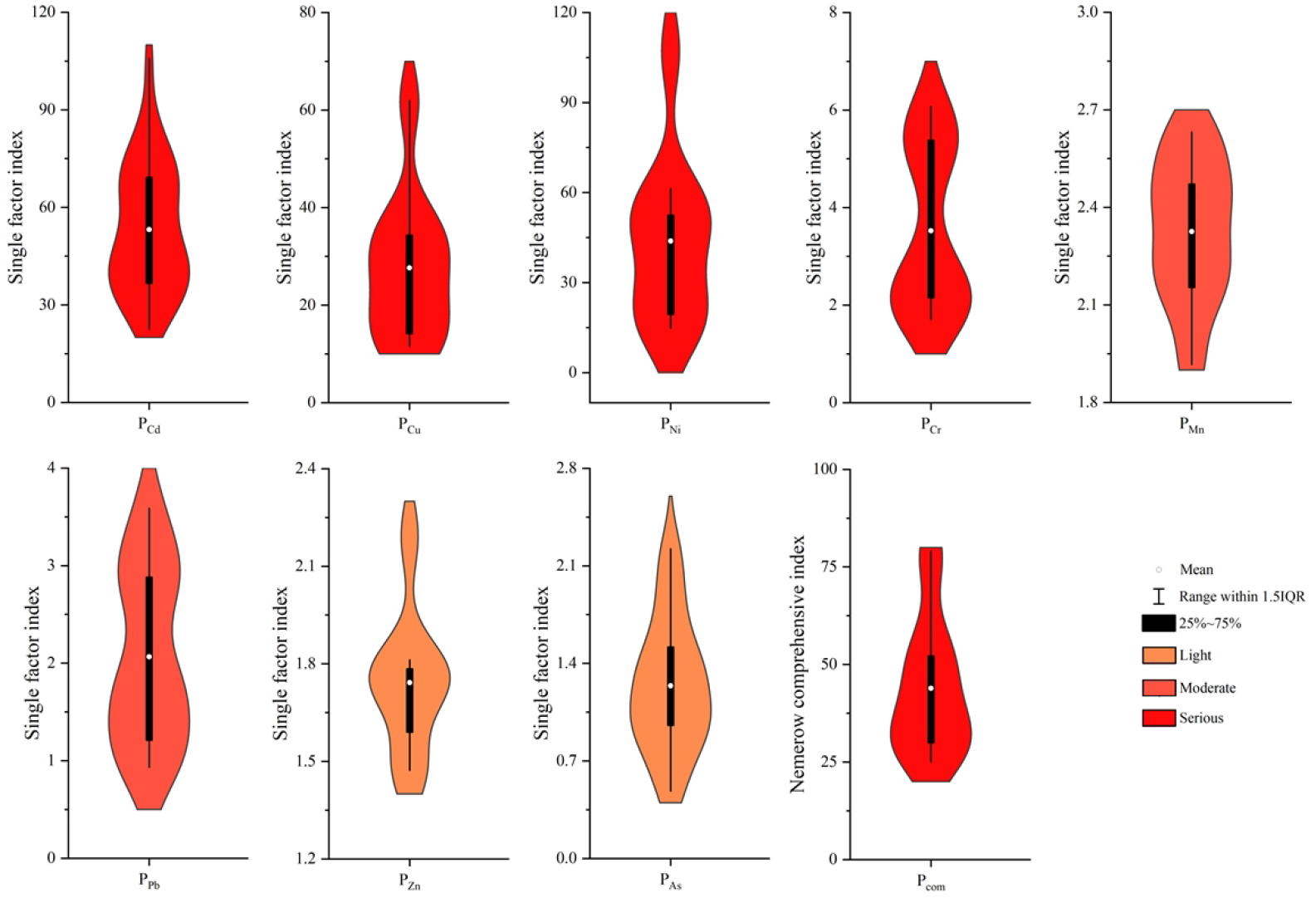
Descriptive statistics for the heavy metal pollution indexes.

### 3.3. Source analysis of heavy metals in soil

Correlation analyses and the positive matrix factorization (PMF) model can be used to analyze the sources of heavy metals in soil (Huang et al., 2021). The Pearson correlation analysis of the heavy metal contents in the soils showed that there were correlations among most of the heavy metals (Fig. 3). Among them, the correlations between Ni and Cu, Zn, Pb, Cr and Cd, Pb, Cu, and Zn, and Pb, Cd, and Pb reached a very significant level (P < 0.01). These results showed that there were strong correlations among some of the heavy metals. The correlations between Ni and Cd, and Cu and Cd, Zn, and Pb reached significant levels (P < 0.05). The analysis results for the sources of heavy metals in the soil obtained from the study area using the PMF model (Fig. 3) show that Factor 1 contributed most to Cd and Cr levels at 44.6% and 42.8%, respectively. Atmospheric deposition is an important source of Cr and Cd in soil. The annual input of Cd and Cr into soil through atmospheric deposition is 493 and 7392 t/a, respectively (Luo et al., 2009). In addition, Cd is also considered to be a symbol element of agricultural activities because chemical fertilizers, livestock manure, and irrigation water are important sources of Cd (Cai et al., 2019; Pan et al., 2016). It has also been reported that the Cd inputs due to fertilizer, livestock manure, and irrigation water into agricultural soil in China are 113, 778, and 30 t/a, respectively (Luo et al., 2009). The study area was located in the arid area of Northwest China, which uses large amounts of water for agricultural irrigation and has developed animal husbandry. These activities may have led to the Cd enrichment seen in the study soil. Therefore, Factor 1 came from composite pollution sources, including atmospheric deposition and agricultural pollution sources. Factor 2 contributed most to Cu and Ni levels at 62.5% and 66.5%, respectively. The correlation analysis showed that the correlation between Cu and Ni was high at 0.88, indicating that Cu and Ni had the same pollution source. The sampling points were located near the copper nickel mining, smelting, and tailings pond, which suggests that Factor 2 is based on industrial pollution sources. Factor 3 contributed 54.5%, 47.9%, and 40.0% to Mn, As, and Zn, respectively. Table 1 shows that the variation indexes for Mn, As, and Zn were low; and the single factor pollution evaluation results (Fig. 2) showed that Mn was moderately polluting, whereas As and Zn were slightly polluting. These results consistently showed that Mn, As, and Zn were less disturbed by human factors. Previous studies have shown that Mn mainly came from soil parent materials (Luo et al., 2022; Lv, 2019; Lv et al., 2015). Therefore, Factor 3 was based on natural sources. Factor 4 made the greatest contribution toward Pb levels at 41.2%. Lead is the main marker for traffic emissions caused by fuel combustion, and engine and catalyst use (Guan et al., 2018). It is estimated that automobile exhaust emissions account for about two-thirds of global lead emissions (Cui et al., 2018; Fei et al., 2020). Therefore, Factor 4 was based on traffic pollution sources.

**Figure 3.**
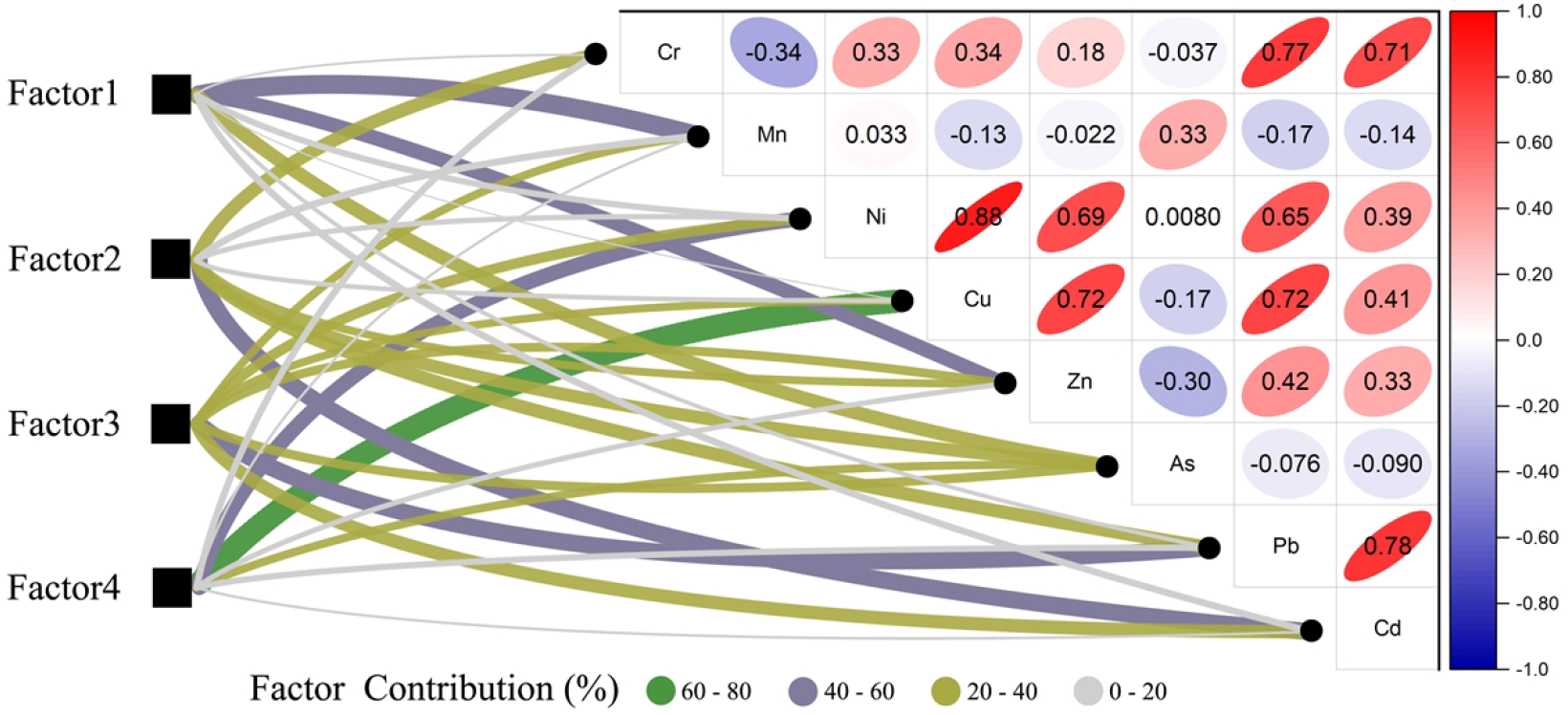
Correlation analysis and source analysis of the heavy metals.

### 3.4. Heavy metal content in plants

The heavy gold contents in different plants around the tailings pond are shown in Fig. 4. It showed that the heavy metal levels in different plants or different parts of the same plant showed large variations, which may be closely related to the physical and chemical properties of the soil, the occurrence forms of heavy metals, and the characteristics of the plants themselves. The contents of eight heavy metals (Cr, Ni, Cu, Zn, Cd, Pb, Mn, and As) in the aboveground parts of 10 plants varied from 2.02–53.77, 24.57–178.67, 7.75–101.67, 10.22–77.37, 6.19–47.13, 0.12–1.72, 0.01–0.94, and 0.28–2.91 mg /kg, respectively, and the Cr, Ni, Cu, Zn, Cd, Pb, Mn, and As variation ranges for their contents in the underground parts of the 10 plants were 5.34–21.50, 26.10–255.33, 21.0–-102.67, 10.22–91.10, 10.7–65.73, 0.17–0.32, 0.03–1.16, and 0.42–5.41 mg/kg, respectively. In general, the Ni, Cu, Pb, and As contents in the underground parts of most plants were higher than those in the aboveground part, while Cr, Mn, and Cd were lower than those in the aboveground parts. The heavy metal contents in plants from high to low were Mn > Ni > Cu > Zn > Cr > Pb > As > Cd. Compared to the maximum values for normal heavy metal contents in plants (Kabata et al., 2007), the heavy metal elements Ni (Ni normal range: 0.1–5 mg/kg) and Cr (Cr normal range: 0.1–0.5 mg/kg) in the underground or aboveground parts of all plants exceeded the maximum value for the normal content range. The Cr content in the aboveground parts of *Atriplex patens* and *Ammophila breviligulata* exceeded the maximum toxicity threshold (toxicity threshold range for Cr: 5–0.5 mg/kg). The Ni content in the underground part of *Klasea centauroides* and the aboveground part of *A. breviligulata* also exceeded the maximum toxicity threshold range (toxicity threshold range for Ni: 10–100 mg/kg). Except *P. australis* and *L. sinense*, the Cu levels in the underground parts of other plants exceeded the maximum normal level (normal range for Cu: 5–30 mg/kg), while the Cu levels in most aboveground parts were within the normal range. The Cd levels in the plants (aboveground or underground parts) exceeded the maximum for the normal range (normal range for Cd: 0.01–0.2 mg/kg), except for *L. sinense*, *S. salsula* and *P. australis*. The As content in all plants (aboveground and underground parts), except for *A. breviligulata*, did not exceed the maximum normal range (normal range for As: 1–1.5 mg/kg), and the Mn, Zn, and Pb contents in all plants (aboveground and underground parts) were within the normal range (normal range for Mn: 30–300 mg/kg, normal range for Zn: 25–250 mg/kg, and normal range for Pb: 5–10 mg/kg).

**Figure 4.**
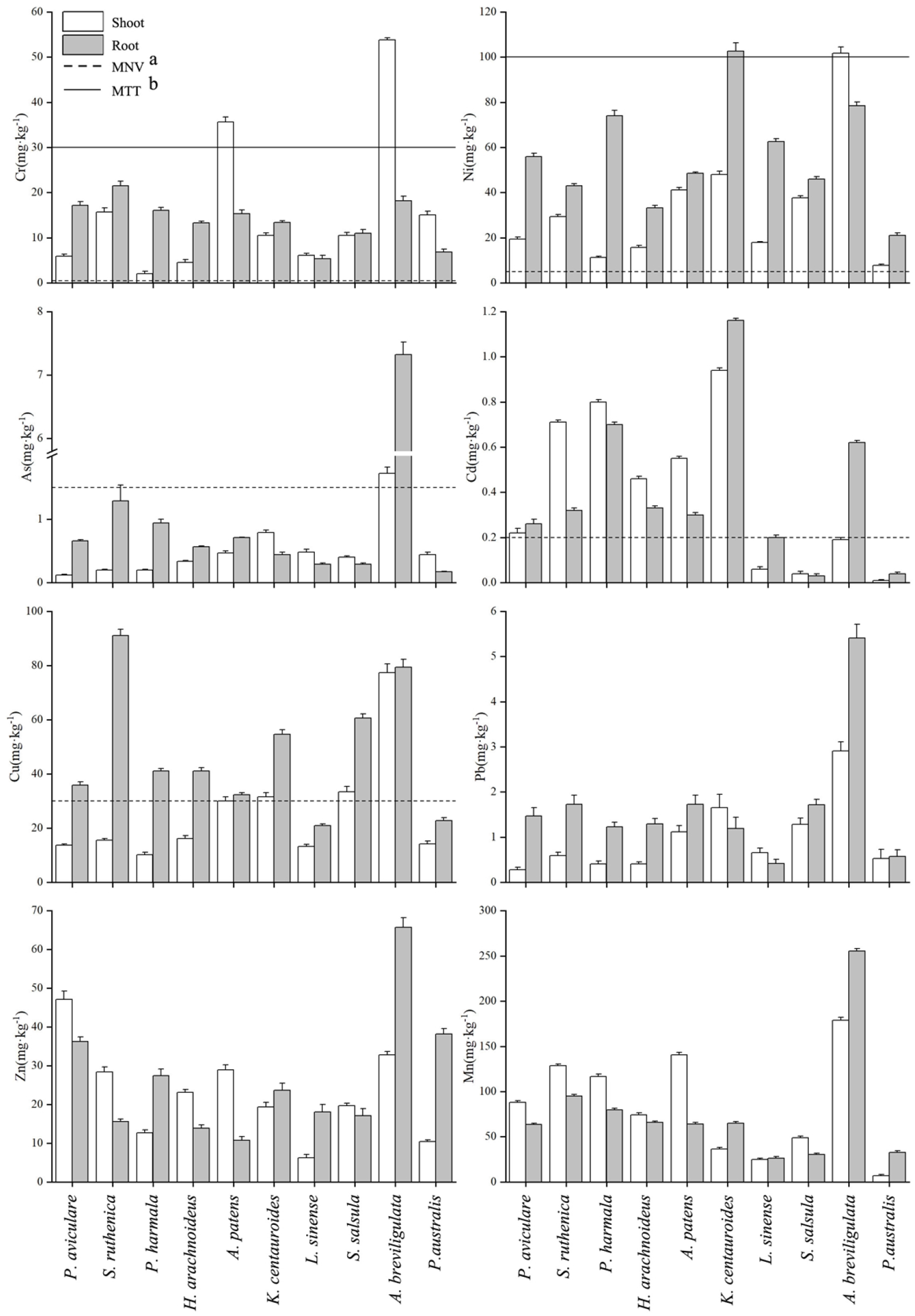
Heavy metal contents in the aboveground and underground parts of 10 desert plants (a: MNV, Maximum normal value, b: MTT, Maximum toxicity threshold).

### 3.5. Heavy metal absorption characteristics of plants

The biological accumulation factor (BAF) shows the ability of plant stems and leaves to accumulate heavy metals, which means that it reflects the ability of plants to remove heavy metals from soil. Table 3 shows that there were obvious differences in the accumulation capacity of plants for heavy metals. It also shows the differences among different plants for the same element and in the same plant for different elements. The minimum biological accumulation factors for the 10 plants around the tailings pond for Cr, Mn, Ni, Cu, Zn, As, Cd, and Pb are 0.006, 0.003, 0.014, 0.051, 0.010, 0.001, and 0.008, respectively, and the maximum values are 0.110, 0.113, 0.104, 0.107, 0.457, 0.133, 0.190, and 0.074, respectively. Among the 10 plants, *A. patens* has a strong ability to extract Cr, *P. aviculare* has a strong ability to extract Zn, *S. ruthenica* has a strong ability to extract Cd, and *A. breviligulata* shows a strong extraction ability for Mn, Ni, Cu, As, and Pb.

**Table 3.** Biological accumulation factors (BAFs) for desert plants around the tailings pond

| Species | BAF |  |  |  |  |  |  |  |
| --- | --- | --- | --- | --- | --- | --- | --- | --- |
|  | Cr | Mn | Ni | Cu | Zn | As | Cd | Pb |
| *P. aviculare* | 0.012 | 0.049 | 0.014 | 0.023 | 0.457 | 0.01 | 0.05 | 0.008 |
| *S. ruthenica* | 0.048 | 0.087 | 0.074 | 0.017 | 0.238 | 0.019 | 0.19 | 0.032 |
| *P. harmala* | 0.006 | 0.08 | 0.022 | 0.032 | 0.084 | 0.035 | 0.176 | 0.02 |
| *H. arachnoideus* | 0.015 | 0.044 | 0.022 | 0.042 | 0.198 | 0.039 | 0.083 | 0.017 |
| *A. patens* | 0.11 | 0.078 | 0.04 | 0.054 | 0.284 | 0.042 | 0.109 | 0.031 |
| *K. centauroides* | 0.014 | 0.023 | 0.03 | 0.039 | 0.158 | 0.049 | 0.097 | 0.029 |
| *L. sinense* | 0.008 | 0.019 | 0.013 | 0.016 | 0.051 | 0.031 | 0.007 | 0.011 |
| *S. salsula* | 0.012 | 0.031 | 0.027 | 0.02 | 0.161 | 0.027 | 0.005 | 0.019 |
| *A. breviligulata* | 0.106 | 0.113 | 0.104 | 0.107 | 0.276 | 0.133 | 0.03 | 0.074 |
| *P. australis* | 0.018 | 0.017 | 0.003 | 0.014 | 0.084 | 0.022 | 0.001 | 0.01 |

Plant stabilization is the use of plant roots to absorb and accumulate toxic metals in soil, thereby reducing the bioavailability and migration of heavy metals in soil (Flathman et al., 1998). The biological concentration factor (BCF) can reflect the ability of plants to stabilize heavy metals in soil. Table 4 shows that the minimum plant stability coefficients for the 10 plants around the tailings pond for Cr, Mn, Ni, Cu, Zn, As, Cd, and Pb were 0.007, 0.019, 0.007, 0.022, 0.105, 0.008, 0.004, and 0.007, respectively, and the maximum values were 0.066, 0.162, 0.143, 0.127, 0.553, 0.567, 0.154, and 0.138, respectively. Among them, *S. ruthenica* can strongly stabilize Cr, *P. harmala* can strong stabilize Ni, Cu, and Cd, and *A. breviligulata* can strongly stabilize Mn, Zn, As, and Pb.

**Table 4.** Biological concentration factors (BCFs) for desert plants around the tailings pond

| Species | BCF |  |  |  |  |  |  |  |
| --- | --- | --- | --- | --- | --- | --- | --- | --- |
|  | Cr | Mn | Ni | Cu | Zn | As | Cd | Pb |
| *P. aviculare* | 0.036 | 0.035 | 0.04 | 0.06 | 0.351 | 0.056 | 0.059 | 0.043 |
| *S. ruthenica* | 0.066 | 0.064 | 0.109 | 0.099 | 0.131 | 0.121 | 0.086 | 0.094 |
| *P. harmala* | 0.048 | 0.055 | 0.143 | 0.127 | 0.182 | 0.164 | 0.154 | 0.061 |
| *H. arachnoideus* | 0.045 | 0.039 | 0.047 | 0.108 | 0.118 | 0.065 | 0.06 | 0.055 |
| *A. patens* | 0.047 | 0.036 | 0.047 | 0.059 | 0.105 | 0.064 | 0.06 | 0.047 |
| *K. centauroides* | 0.019 | 0.042 | 0.063 | 0.067 | 0.193 | 0.027 | 0.119 | 0.021 |
| *L. sinense* | 0.007 | 0.02 | 0.046 | 0.026 | 0.15 | 0.018 | 0.022 | 0.007 |
| *S. salsula* | 0.012 | 0.019 | 0.033 | 0.037 | 0.141 | 0.02 | 0.004 | 0.025 |
| *A. breviligulata* | 0.036 | 0.162 | 0.081 | 0.109 | 0.553 | 0.567 | 0.097 | 0.138 |
| *P. australis* | 0.008 | 0.019 | 0.007 | 0.022 | 0.309 | 0.008 | 0.005 | 0.011 |

### 3.6. Comprehensive evaluation of phytoremediation potential

The plant comprehensive extraction index and plant comprehensive stability index results, calculated based on Fuzzy evaluation (Fig. 5), showed that among the 10 plants investigated, the plant comprehensive extraction indexes for *P. australis*, *L. sinense*, *S. salsula*, *H. arachnoideus*, *K. centauroide*s, *P. harmala*, and *S. ruthenica* were 0.04, 0.05, 0.14, 0.23, 0.25, 0.26, and 0.30, respectively, which were less than 0.4, which meant that their comprehensive removal potentials for heavy metals were poor. The plant comprehensive extraction index for *A. patens* was 0.52, which meant that its comprehensive removal potential for heavy metals was good; and the plant comprehensive extraction index for *A. breviligulata* was 0.83, which means that its comprehensive removal potential level for heavy metals was excellent, indicating that it had a strong comprehensive removal ability for heavy metals. The plant comprehensive stability indexes for *P. australis*, *L. sinense*, *S. salsula*, *A. patens*, *K. centauroide*s, *P. aviculare*, and *H. arachnoideus* were smaller at 0.07, 0.07, 0.08, 0.28, 0.29, 0.31, and 0.34, respectively, which are all less than 0.4. Therefore, their potential stability indexes for heavy metals are poor, indicating that their comprehensive ability to stabilize heavy metals is weak. The plant comprehensive stability indexes for *S. ruthenica* and *P. harmala* were 0.53 and 0.60 respectively, which meant that their potential stability grades for heavy metals were good. The plant comprehensive stability index for *A. breviligulata* was 0.81, which meant that its stability potential index for heavy metals was excellent, indicating that it had strong comprehensive ability to stabilize heavy metals. Therefore, *A. breviligulata* had greater remediation potential for heavy metals in soil than the other plants.

**Figure 5.**
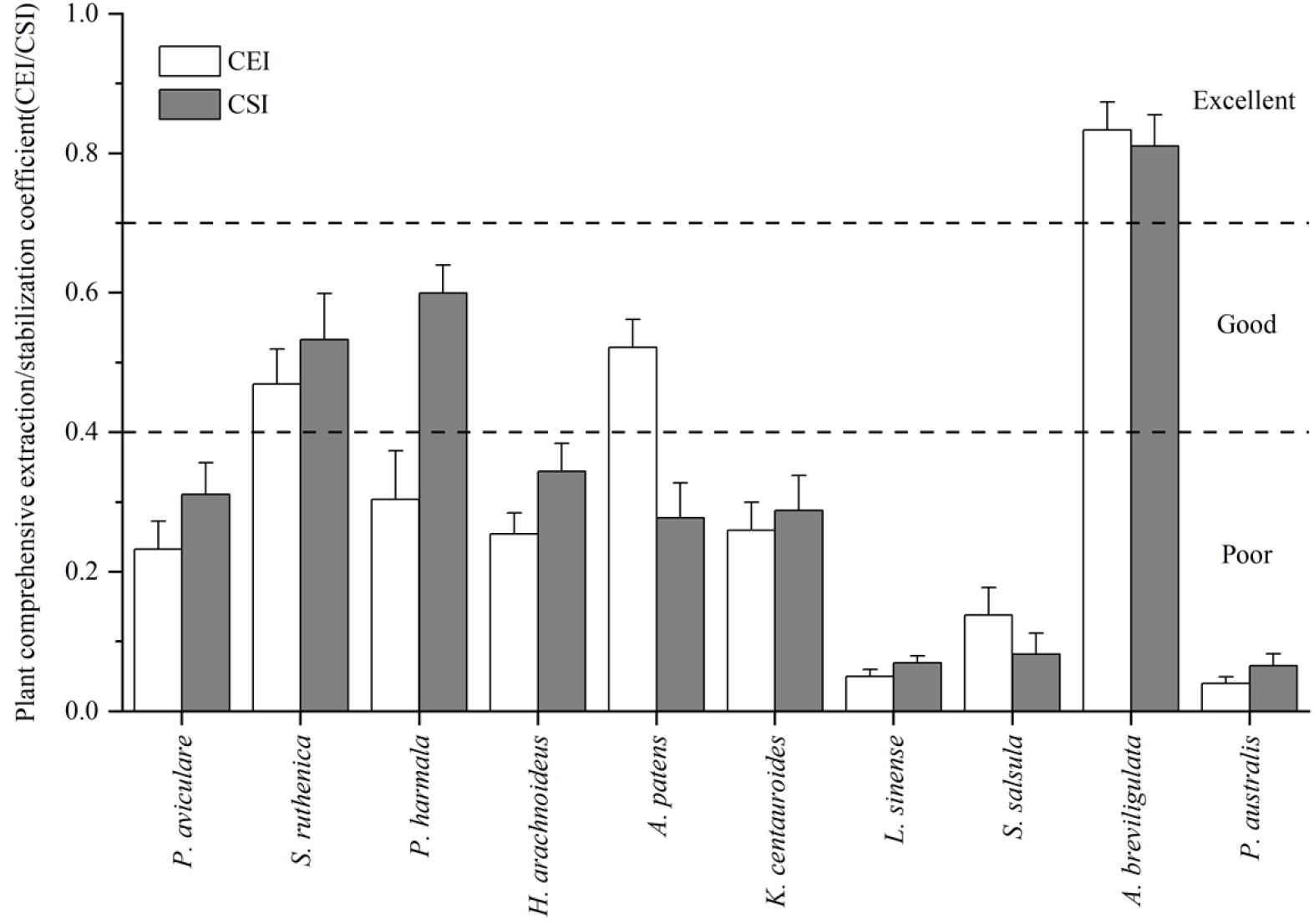
Plant comprehensive enrichment/stability coefficients for the 10 desert plants.

## 4. Discussion

Tailings ponds are one of the important sources of soil heavy metal pollution. Climatic factors (temperature, precipitation, and wind) play an important role in aggravating the migration and diffusion of heavy metal pollution (Meza Figueroa et al., 2009b; Punia, 2021). Temperature affects the oxidation-reduction reaction rate in the environment and the oxidation-reduction reaction affects the mobility of heavy metals. Previous research has shown that, compared to the initial stable heavy metal activities, the oxidation-reduction reaction in mine tailings led to an increase in Pb, Zn, and Mn activities by 42–69% (Trifi et al., 2019). In cold regions, the amounts of heavy metals leached in tailings increases with the increase in temperature (Fu et al., 2019). Kalinovic et al. (2019) investigated the heavy metal contents in soil and found that wind direction had an important impact on the distribution of heavy metal pollution in soil and that the heavy metal contents in the soils that were in the path of the prevailing wind were higher. Tailings have low pHs, water retention capacities, and cation exchange capacities, which means that heavy metals in tailings exhibit strong fluidity (Meza-Figueroa et al., 2009a) and that heavy metals will show accelerated diffusion under high precipitation and snow melt conditions. In arid areas, strong surface evaporation promotes the formation of sulfates on the surface of mining wastes (Khorasanipour et al., 2018) These sulfates are prone to aeolian diffusion and have a negative impact on the surrounding environment (Meza-Figueroa et al., 2009b). The study area was located in the arid area of Northwest China where the temperature difference between day and night is large, it is hot in summer and cold in winter, and there is strong evaporation and erosion (wind erosion, hydraulic erosion, and freeze-thaw erosion). Therefore, there may be a potential risk of heavy metal diffusion in the tailings pond. The evaluation results for heavy metals in the soil around the copper-nickel tailings pond in this study showed that Cd, Cu, Ni, and Cr were at the heavy pollution level, Mn and Pb were at the moderate pollution level, and Zn and As were at the light pollution level. The source analysis using the PMF model showed that pollution by Cu, Ni, Cr, and Cd was closely related to industrial activities, such as mining, ore smelting, and tailings accumulation. Li et al. (2022) found that there were many kinds of heavy metal pollution in the soil around a copper nickel mine tailings pond, among which Cu, Ni, and Cr were the main environmental pollutants derived from tailings. Their results were similar to the results obtained from this study. Liang et al. (2017) collected water, soil, rice, and vegetable samples in an area near a tailings pond and found that these samples were contaminated with heavy metals and posed potential health risks to nearby residents. Therefore, the diffusion of heavy metals in tailings ponds will cause serious pollution to the environment and pose a serious threat to human life and health.

Phytoremediation uses the accumulation of elements by plant stems, leaves, and roots to remove and stabilize heavy metals in soil (Fig. 6). Furthermore, the plant canopy formed after phytoremediation can reduce the near surface wind speed and the diffusion of fine particle pollutants, while the underground root network can prevent rainfall erosion and leaching, and provide a suitable rhizosphere environment for heavy metal precipitation (Cameselle et al., 2013; Houben et al., 2013). In general, identifying the natural vegetation growing on contaminated sites and the selection of metal tolerant plants with potential remediation value are some of the most important components of a localized phytoremediation strategy (Matanzas et al., 2021). Mousavi et al. (2020) evaluated the remediation potential of plant species growing in a heavy metal polluted saline alkali soil and found that seven plants (*L. arborescens*, *A. santolina*, *P. gnaphalodes*, *Z. eurypteru*, *P. harmala*, *P*. *olivieri*, and *A. javanica*) have good heavy metal stabilization effects, while *Z. eurypterum* and *A. javanica* also have good heavy metal and relatively high heavy metal extraction abilities, respectively. Liu et al. (2016) evaluated the plant stability and plant extraction potential of desert plants naturally growing around tailings ponds in Altay, Xinjiang Province, China. The evaluation results showed that *S. schmidt* was most suitable for stabilizing Cu, *K. caspica* was most suitable for stabilizing Cd, and *P. Aviculare* had a high extraction potential for Cu. Zhao et al. (2014) used a fuzzy comprehensive evaluation to evaluate the comprehensive heavy metal extraction potential of woody plants growing on heavy metal contaminated sites and found that *B. papyrifera* could extract many kinds of heavy metals at the same time. In terms of extraction potentials for single heavy metals, this study found that *A. patens* had a strong ability to extract Cr; *P. aviculare* had a strong Zn extraction ability; *S. ruthenica* had a strong Cd extraction ability; *A. breviligulata* had strong Mn, Ni, Cu, As, and Pb extraction abilities. In terms of single heavy metal extraction and stabilization abilities, this study found that *S. ruthenica* had a strong ability to stabilize Cr; *P. harmala* had a strong ability to stabilize Ni, Cu, and Cd; and *A. breviligulata* had a strong ability to stabilize Mn, Zn, As, and Pb. The evaluation results for plant heavy metal comprehensive extraction/stabilization potential based on fuzzy mathematics showed that the plant comprehensive extraction and plant comprehensive stability indexes for *A. breviligulata* were the highest, indicating that *A. breviligulata* had better heavy metal removal and stabilization effects than the other native plants. Therefore, *A. breviligulata* can be selected as the preferred species for heavy metal pollution remediation in the study area.

**Figure 6.**
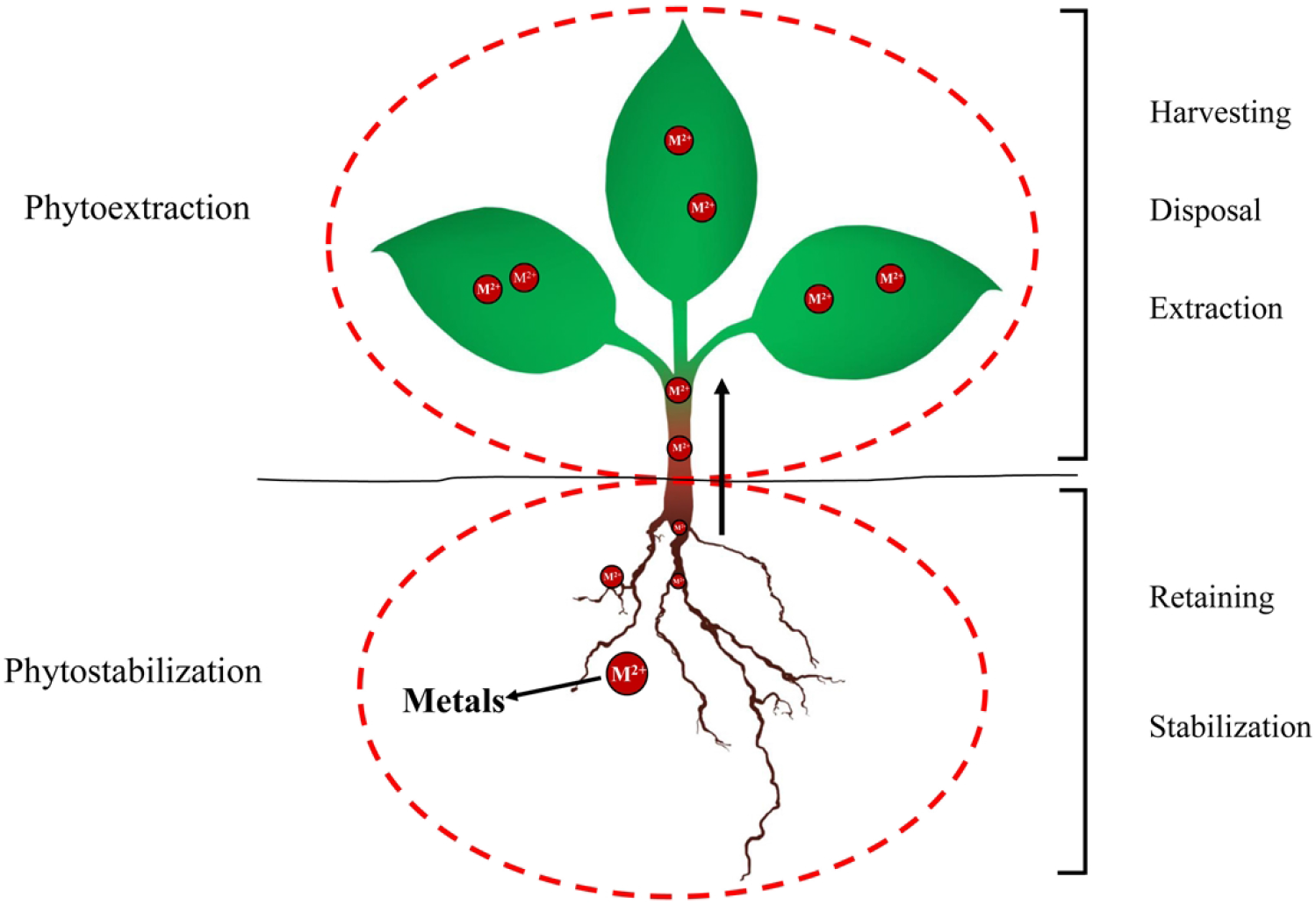
Phytoextraction and phytostabilization.

## 5. Conclusion

The results showed that there was obvious compound heavy metal pollution in the soil around the copper-nickel tailings reservoir area. Among them, the heavy metal elements Cd, Cu, Ni, and Cr were at the heavy pollution level; Mn and Pb were at the moderate pollution level; and Zn and As were at the slight pollution level. The PMF model results showed that the Cu and Ni came from industrial pollution sources; Cd and Cr came from atmospheric deposition and agricultural pollution sources; Pb came from traffic pollution sources; and Mn, Zn, and As came from natural sources. Compared to the normal heavy metal content in plants, the heavy metal elements in the underground and aboveground parts of the 10 desert plants surveyed exceeded the standard, indicating that heavy metal pollution in the soil posed a threat to plant growth. Among the 10 desert plants, the CEI and the CSI values for *A. breviligulata* were highest, indicating that it has a good potential to remove and stabilize a variety of heavy metals in the soil. Therefore, *A. breviligulata* had a strong comprehensive remediation potential for heavy metals in the soil around the tailings pond and it could be used to remediate heavy metal pollution in the soil. This study used fuzzy evaluation to evaluate the comprehensive extraction/stability potential of plants at a site with multiple heavy metal compound pollution. The processes used in this study successfully identified local plants with higher polymetallic remediation capacities that could be utilized in local remediation projects.

## Data availability

All relevant data are within the manuscript and its Supporting Information files.

## Author contribution

Conceptualization: Jianfei Shi, Zhengzhong Jin, Zhibin Zhou.

Data curation: Jianfei Shi, Wenting Qian.

Formal analysis: Jianfei Shi.

Investigation: Jianfei Shi, Xin Wang, Xiaoliang Yang.

Methodology: Jianfei Shi.

Supervision: Jianfei Shi, Zhengzhong Jin.

Writing-Original draft: Jianfei Shi.

Writing-review & editing: Jianfei Shi, Zhengzhong Jin, Zhibin Zhou.

